# Identification of host antiviral genes differentially induced by clinically diverse strains of Tick-Borne Encephalitis Virus

**DOI:** 10.1101/2021.02.23.432617

**Authors:** Niluka Goonawardane, Laura Upstone, Mark Harris, Ian M Jones

## Abstract

Tick Borne Encephalitis Virus (TBEV) is an important human arthropod-borne virus, which causes tick-borne encephalitis (TBE), an acute viral infection of the central nervous system (CNS) that causes neurological symptoms of varying severity. TBEV is prevalent in large parts of central- and northern-Europe as well as Northern Asia, and strains of varying pathogenicity have been described. Both host and viral specific characteristics have been postulated to determine the outcome of TBEV infection, but the exact basis of their clinical variability remains undefined.

Here, we report the generation of Spinach RNA aptamer labelled TBEV replicons of high (Hypr) and low (Vs) pathogenicity isolates and perform the first direct comparison of both strains in cell culture. We show that pathogenic Hypr replicates to higher levels than Vs in mammalian cells, but not in arthropod cells, and that the basis of this difference maps to the NS5 region, encoding the methyltransferase and RNA polymerase. For both Hypr and Vs strains, NS5 and the viral genome localized to defined intracellular structures typical of positive strand RNA viruses, but Hypr was associated with significant activation of IRF-3, caspase-3 and caspase-8, whilst Vs activated Akt, affording protection against caspase-mediated apoptosis. Activation of TIAR and the formation of cytoplasmic stress granules were an additional early feature of Vs but not Hypr replication. Taken together, these findings highlight NS5 and novel host cell responses as key underling factors for the differential clinical characteristics of TBEV strains.

**Importance:** Tick-borne encephalitis virus (TBEV) is an emerging virus of the flavivirus family spread by ticks. Tick bite can transfer the virus and lead to a febrile infection, Tick-borne encephalitis, of varying severity. There is no specific therapeutic treatment and control in endemic areas is by vaccination. The basis of the different pathologies shown following TBEV infection, from mild to fatal, is not clear although the virus genotype clearly has a role. Mapping the basis of their differential effects would allow focus on the stages of the replication cycle responsible, which might guide the development of therapeutic interventions or the creation of purposefully attenuated strains as candidate vaccines.

## Introduction

Tick-borne encephalitis virus (genus Flavivirus, family *Flaviviridae)* is the causative agent of Tick-borne encephalitis (1), an important arthropod borne disease of the central nervous system (CNS) that is endemic in parts of Europe and Asia (2). Now recognised as a re-emerged human pathogen, TBEV has spread into new geographical areas, with an estimated 14,000 TBEV cases reported across 30 European and Asian countries annually (1, 3, 4). In 2019, TBEV was isolated from ticks in the East of England, highlighting its potential emergence in the UK (5). Three closely related groups of TBEV exist, European (TBEV-Eur), Far Eastern (TBEV-FE) and Siberian (TBEV-Sib), although more recent sequence-based analysis suggests further distinct sub-divisions of the Siberian lineages (6). Although genetically similar, the clinical outcome of infection by different TBEV strains is variable, ranging from asymptomatic in 80-98% of cases to encephalitis of varying severity (2, 3, 7), but the basis of this diversity remains unclear. In nature, TBEV is maintained by *Ixodes* ticks, with the majority of human TBEV infections caused by bites from *I.ricinus* (European and Far Eastern strains) or *I.persulcatus* (TBEV-Sib strain) (8, 9).

TBEV is an icosahedral enveloped (~50 nm) virus with a positive-sense RNA genome of ~11 kb flanked by 5- and 3-prime untranslated regions (UTRs) (10). The capped viral RNA is translated into a single polyprotein precursor that is cleaved by cellular and viral proteases to yield three structural proteins (C, prM, and E) and seven non-structural proteins (NS1, NS2A, NS2B, NS3, NS4A, NS4B and NS5) (11, 12). The C-terminal domain of NS5 encodes the viral RNA-dependent RNA polymerase (RdRp) (13). In common with other positive sense RNA viruses, TBEV induces host cell membrane rearrangements to establish compartmentalised viral factories (14, 15), postulated to protect viral RNAs from host defences (15). TBEV infection triggers innate immune signalling through its interaction with RIG-I/MDA5, which promotes IRF-3 translocation to the nucleus (16, 17). Specific antiviral response genes have recently been shown to supress TBEV replication in culture, including TRIM79α in mouse cells (18) and TRIM5α (19) and viperin (20) in human cells. The chemokine RANTES is upregulated in response to TBEV infection in human neuronal tissue, via IRF-3 activation, and has been linked to TBEV neuropathology (17). TBEV also induces caspase-3 dependent apoptosis (21), which can be delayed by the interferon antagonist function of NS5 (22–25) and the NS4A mediated inhibition of STAT signalling (26). These mechanisms (summarized in (27)) are strain-specific and could contribute to the variable clinical pathology of different TBEV isolates.

TBEV strain Vasilchenko (Vs) was first isolated in the Novosibirsk region of Russia in 1969 (28) and infectious clones have been generated (29). Vs causes a subclinical infection *in vivo* and shows minimal cytopathic effect (CPE) in cell culture (29). In contrast, strain Hypr, isolated in 1953 from an infected Czechoslovakian child, exhibits extensive CPE in cell culture and significant neuro-invasiveness in mice (30). Despite these differences, Vs and Hypr strains are ~96% homologous at the amino acid level, permitting gene exchange studies to identify loci related to their pathogenicity. In this regard, Khasnatinov *et al*., developed Vs/Hypr chimeras revealing that the non-viremic transmission (NVT) of TBEV amongst ticks co-feeding on mice (31) is mediated by the 5-prime region of the genome encoding the structural E protein. In contrast, CPE was mapped to the 3-prime region encoding the non-structural proteins but it was not investigated in further detail (Khasnatinov *eŕ al., ibid*) and the clinical variability of TBEV remains largely undefined.

Here, to address this issue, we generated a novel series of Hypr and Vs replicons and chimeras in which the structural genes were replaced with a *cis* acting RNA tag Spinach2 (32) to directly visualize RNA and the kinetics of virus replication. Chimeras in which the NS3 or NS5 regions of the Hypr/Vs were exchanged were further generated for a comparison of their replication characteristics in both mammalian and insect cell lines. The induction of early host innate immune defence, including IRF-3, caspase-3, caspase-8, phospho-Akt, and TIAR were also measured between Vs/Hypr hybrids to compare their ability to regulate host cell defences. We herein reveal key viral and cellular determinants that may contribute to the variable virus neuropathology of Hypr and Vs strains.

## Materials and Methods

### Cells

Porcine embryo kidney (PS) cells were grown in RPMI 1640 medium (Gibco^®^ Life technologies UK) supplemented with 5% foetal bovine serum (FBS) at 37°C in a CO2 (5%) incubator. Unless otherwise stated, replicon transfected PS cells were maintained medium supplemented with in a 2% FBS and 1% Penicillin-Streptomycin (Invitrogen). Sf9 cells (ATCC) were cultured in BioWhittaker Insect-Xpress supplemented with 2% FCS, 1% Penicillin-Streptomycin and 2.5 μg/ml amphotericin B. Cells were grown at 28°C as monolayers or in suspension with agitation at 100 rpm. Transfections of TBEV replicons were performed in monolayer cultures of Sf9 cells.

### Construction of TBEV replicons

In this study we used cDNA clones of WE-TBEV strain Hypr (U39292) or SIB-TBEV strain Vs (AF069066) (29, 31) positioned downstream of the SP6 promoter, from which infectious RNA could be transcribed *in vitro*. Chimeric replicons were constructed through restriction fragment swapping, using intermediate plasmids as required, or through synthesis of DNA fragments *de novo* followed by Gibson assembly (33) using NEBuilder HiFi (NEB). All clones were sequenced throughout the TBEV genome for verification.

### Recovery of replicon RNA

DNA plasmids encoding the relevant clone or hybrid were linearised at the *SmaI* restriction site downstream of the TBEV coding sequence and used as template to produce full-length capped RNA using SP6 RNA polymerase (Promega USA). Each reaction (50 μl total volume) contained 1 μg of linearised DNA template and 40 units of SP6 RNA polymerase, incubated at 37°C for 3 h. RNAs were purified using the SV total RNA isolation kit (Promega USA) and resuspended in RNase-free water (Invitrogen USA). RNA integrity was verified by agarose gel (1%) electrophoresis and quantified by spectrophotometry.

### Titration of infectious TBEV by plaque assay

TBEV full-length RNAs were produced and purified as described above. PS cells were transfected with the SP6-transcribed RNAs (1 μg RNA for 1.2 x10^5^ cells) using Lipofectamine 2000 (Invitrogen, USA) as per the manufacturer’s recommendation. Infectious supernatant medium was collected at 24 to 72 hpi. Aliquots of virus, diluted with serum-free RPMI 1640 medium, were applied to monolayers of PS cells for 1 h at 37 ^0^C. The inoculum was aspirated, and plates were overlaid with RPMI 1640 medium supplemented with 2% FCS and 1% SeaPlaque Agarose (Cambrex, USA) for 5 days at 37 ^0^C for plaque formation. Monolayers were then fixed with 4% formaldehyde and stained with 0.05% crystal violet. Plaques were counted and virus titre was expressed as log_10_ pfu/ml. All virus work was performed in a Biological Safety Level 3 (BSL3) laboratory. For TBEV growth kinetics, PS cells were infected with WT or NS3/NS5 chimeras at an MOI of 0.1 for 1 h. Infected cells were washed five times with PBS and incubated with fresh complete medium (2% FCS) at 37°C. The supernatant medium from infected cells was collected at 0, 4, 8, 12, 16, 20, 24 and 72 hpi and frozen at –80°C prior to further analysis. The experiments were performed in triplicate. The titres of infectious virus at different time points were determined by plaque assay.

### RNA transfections

Transfections were performed using Lipofectamine 2000 (Invitrogen, USA) as per the manufacturer’s recommendation. Briefly, 1 μg of SP6 transcribed RNA was complexed with the transfection reagent in serum- and antibiotic-free culture medium for 10 min at room temperature, complexes were added to cell monolayers for 24 to 96 h. All transfections were performed in triplicate.

### Immunofluorescence (IF) assays

For IF analysis, transfected cells cultured in 35 mm glass-bottomed dishes (MatTek Corp.), were fixed with 4% paraformaldehyde (Applichem GmbH, Germany) and permeabilised in PBS-T (0.1% v/v Triton X-100 in PBS) for 5 min. Cells were blocked in PBS-T containing 5% w/v BSA for 10 min and probed with primary antibodies in PBS-T, 1% w/v BSA for 1 h at room temperature. After washing with PBS, cells were stained with 1:500 fluorochrome-conjugated secondary antibodies in PBS-T, 1% w/v BSA for 1 h at room temperature in the dark. Nuclei were counterstained with Hoechst 33342 DNA dye NucBlue^®^ Live ReadyProbes^®^ reagent for live cell imaging or 4’,6’-diamidino-2-phenylindole dihydrochloride (DAPI) and SYTO^®^ 60 fluorescent nucleic acid stain for fixed samples (Invitrogen, Molecular Probes, Germany).

### Detection of Spinach2 tagged replicon RNA

For the live-cell analysis of Spinach2 expression, cells were imaged 24-96 h post-transfection. Cell media was replaced with imaging media (RPMI 1640 without phenol red or vitamins, supplemented with 25 mM HEPES, 5 mM MgSO4), 20 μM DFHBI (Lucerna technologies USA) 30 min prior to analysis. Cell images were analysed using NIS-Elements software. Background intensities were subtracted from all pixel intensity measurements to avoid noise in the final images.

### Luciferase assays

PS cells were seeded at 1.5 × 10^5^ cells per well in 96-well plates (n=3) and transfected with *in vitro* transcribed TBEV replicon RNA (100 ng/well) and 10 ng of pTK-Ren (Promega) using Lipofectamine-2000 (ThermoFisher Scientific). Cells were harvested in passive lysis buffer (Promega) from 12-72 hpt and luciferase activity was measured using the Dual Luciferase reagent kit and GloMax multi detection system (Promega). Firefly luciferase readings from the TBEV replicons were normalized to the Renilla values (pTK-Ren) of each sample.

### Confocal microscopy and colocalization analysis

Live fluorescence images were obtained on an A1R laser scanning microscope (LSCM) using an A1^+^’s galvano scanner and oil immersion objective (Nikon, Japan). Spinach2 was imaged using a FITC 488 nm laser for EGPF (470/40 excitation and 515/30 emission). Nuclei were stained with NucBlue^®^ Live Ready Probes^®^ (Invitrogen, USA) and detected using DAPI filters (emission filter 450/35nm). NS5 protein and SGs images were obtained using a Leica TCS SP8 confocal microscope with a 592 nm and/or a 660 nm depletion laser and a HCX Plan Apo 100×/1.4 oil objective. Images were analysed using the FIJI package of Image J. For quantification of the spatial distribution of Spinach2-RNA, images were acquired under identical parameters, but with a variable gain to ensure correct exposure. Two-dimensional areas and foci counts of the aptamer tagged RNAs were measured using the Analyze Particles function in Fiji. Statistical significance was determined using a two-tailed Student’s t-test with Welch’s correction. For co-localisation analysis, Manders’ overlap coefficients were calculated using Fuji software with Just Another Co-localisation Plugin (JACoP) (National Institutes of Health) (34), where the M1 coefficient reports the fraction of the dsRNA signal that overlaps the anti-G3BP1 signal, whilst the M2 coefficient reports the fraction of the anti-G3BP1 signal that overlaps with the dsRNA signal. Coefficient values ranging from 0 to 1, corresponding to non-overlapping images and 100% co-localization, respectively. Co-localisation was assessed on ≥ 10 cells from at least two independent experiments.

### RT-PCR analysis

RT-PCRs were performed using a one-step protocol in a total reaction volume of 25 μL. Reaction mixture contained 5x Qiagen OneStep RT-PCR Buffer, dNTP Mix (containing 10 mM of each dNTP), Qiagen OneStep RT-PCR Enzyme mix with HotStarTaq DNA Polymerase, RNase inhibitor (10 unit), 0.6 μM forward and reverse primers and 1 μg of template RNA. RT reactions were performed at 50°C for 30 min. PCR parameters were as follows: 30 cycles at 94°C for 1 min for denaturation; 65°C for 1 min for primer annealing, and 72°C for 1 min for extension. PCR products were confirmed by electrophoresis in 1% w/v agarose gels.

### Statistical analysis

Statistical significance was determined in GraphPad Prism using a Student’s *t* test with Welch’s correction or a one-way ANOVA with Bonferroni’s correction.

## Results

### Generation of the Spinach TBEV replicon system

Flavivirus replicons have been previously described in which the structural genes are replaced with sequences encoding GFP or luciferase reporters (reviewed in (35)), including for TBEV (36). Upon transfection both the markers and non-structural proteins of the replicons are translated, with the latter recognising the 5’ and 3’ UTRs of the genome to establish active replication within cells. In this study, Hypr and Vs replicons were engineered to express a Spinach aptamer, which forms an RNA sequence that folds to produce a fluorescent signal in the presence of 3,5-difluoro-4-hydroxybenzylidene imidazolinone (DFHBI). Replicons of Hypr and Vs were produced as surrogates for studies on infectious TBEV clones, as fully infectious virus systems are hazard group 3 viruses which require the high level of safety containment, rendering mutational/chimeric analysis unfeasible. The Spinach system has been used to visualize RNA replication in several systems but has not been previously reported for TBEV (37, 38). The Spinach2 aptamer within a tRNA scaffold was incorporated into the +ve strand at the 5’-end of the sequence encoding the C-protein (at codon 41) to maintain the essential cyclization sequence (Figure 1A). The Firefly luciferase gene was cloned downstream of the Spinach2 sequence to provide a marker of TBEV translation. The firefly sequence was modified to contain no CpGs and a low UpA frequency (luc-cu) to enhance the replication capacity as previously described (39). To ensure correct polyprotein processing, the self-cleaving 2A peptide from Foot and Mouth Disease Virus (FMDV) was incorporated downstream of the luc-cu sequence prior to the transmembrane (TM) region of E (22 codons), to ensure correct translocation of the downstream NS1 protein into ER (40) (Figure 1A). The ability of the constructed RNA to produce fluorescence was assessed following SP6 mediated transcription and incubation of the purified transcript with 10 μM DFHBI (Lucerna Technologies USA), followed by and irradiation at 480 nm (Figure 1B).

**Figure 1.**
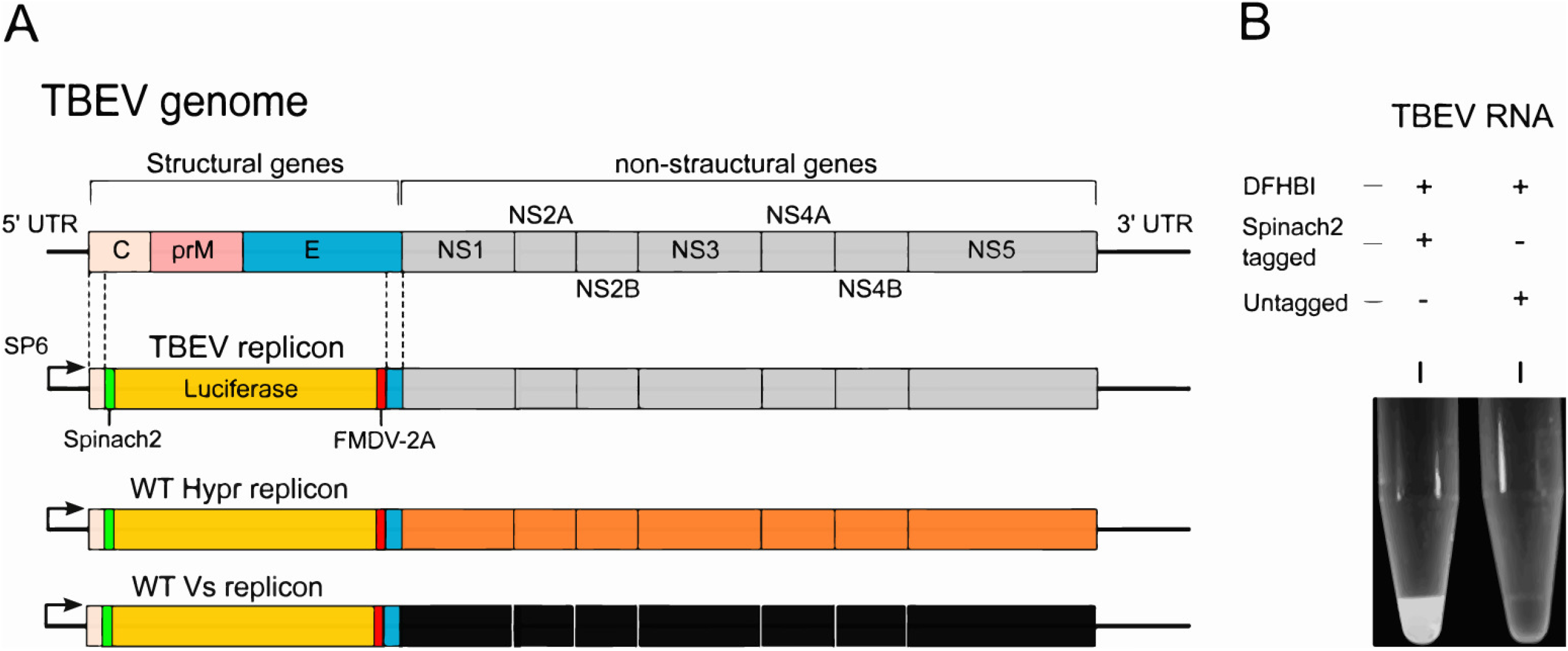
TBEV replicon system. (A) Schematic representation of the TBEV genome (top) and TBEV replicon in which the structural genes were replaced with Spinach2, FMDV-2A or and firefly (cu) luciferase. WT Hypr and Vs replicons are shown as a reference for subsequent hybrid construction. (B) RNA-fluorophore complexes from SP6 transcribed TBEV RNA (1 μg) mixed with the Spinach substrate DFHBI for 30 min.

### Characterisation of the Spinach TBEV replicon system

To validate each genome assembly as a functional replicon, capped RNAs were transfected into porcine kidney stable (PS) cells, chosen as they have previously been shown to support active TBEV replication (41). Cells were harvested at defined time-points post transfection (hpt) and viral genomes were quantified using negative strand RT-PCR (42, 43). Upon analysis, both Hypr and Vs replicons showed evidence of negative strand synthesis from 12 hpt (Figure 2A). Upon assessment of the kinetics of viral RNA synthesis, the Hypr replicon showed robust replicative activity at early time points, peaking at 24 hpt and declined thereafter. In contrast, the Vs replicon produced lower and more gradual levels of RNA synthesis that persisted over a longer time period than the Hypr replicon (Figure 2A). Importantly, these features of Hypr and Vs were not restricted to the replicon system and were confirmed in fully infectious virus systems using capped, *in vitro* transcribed chimeric virus RNA transfected into PS cells, with virus production assessed by plaque assay of the cell supernatants at 24 hours post infection (hpi) (Figure 2B). The Hypr viruses showed more rapid virus production up to 24 hpi compared to the slower more sustained growth of Vs, validating the kinetics of the Spinach TBEV replicon system (Figure 2B). To ensure completion of the Spinach TBEV replication cycle, Firefly luciferase signals, translated from progeny positive sense replicon RNAs were measured in both PS and A549 (alveolar basal epithelial) cells, the latter included as a representative human cell that is known to support flavivirus replication (52). Replicon-derived luciferase activity when normalised to the levels of the co-transfected Renilla luciferase reflected the levels of negative sense RNA, with an early but short-lived peak for Hypr and a slower more prolonged signal for Vs (Figure 2C-D). Of note, the rates of replication for Hypr and Vs were comparable between PS and A549 cells, suggestive of a conserved replicative phenotype across mammalian cell lines.

**Figure 2:**
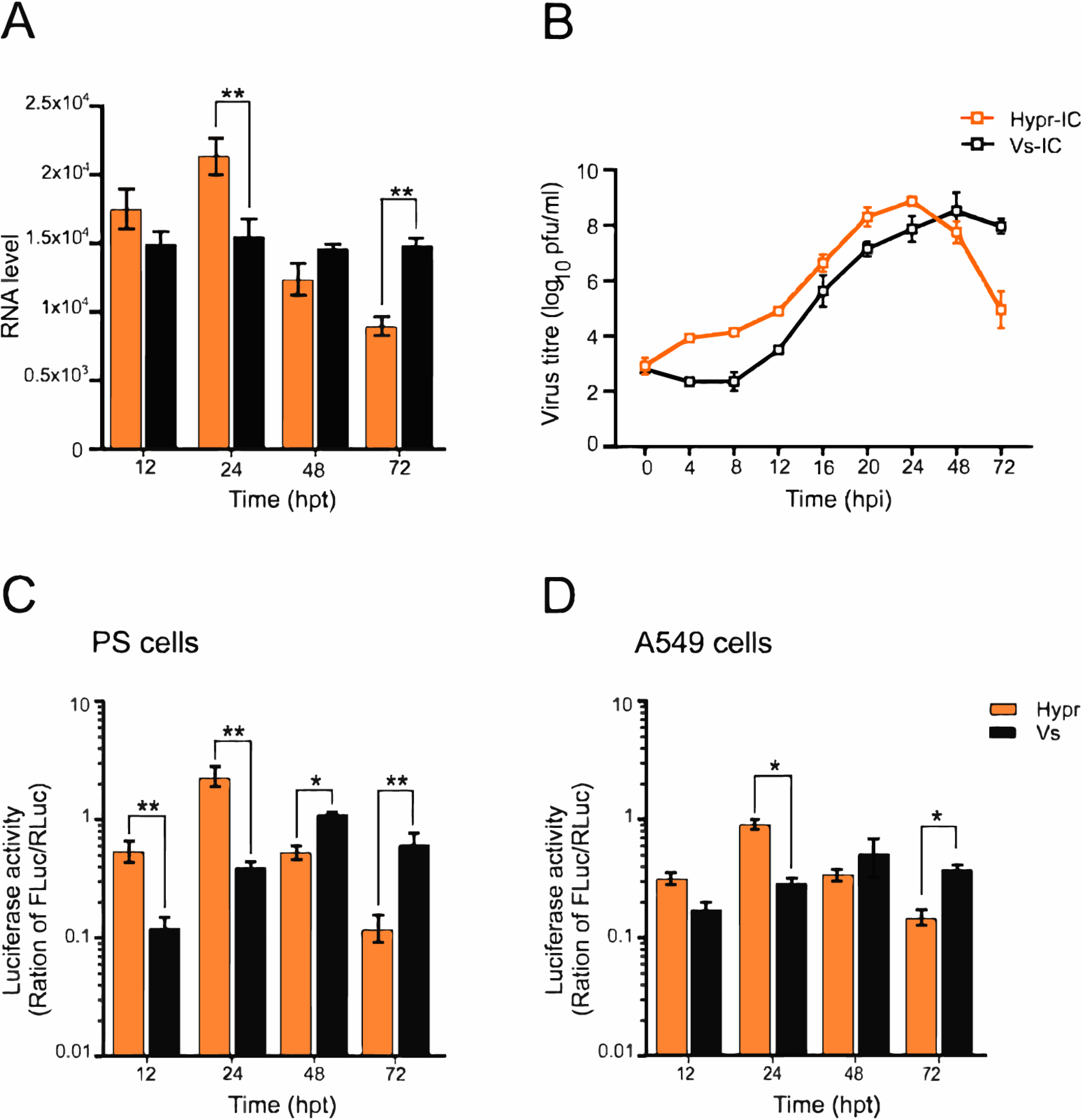
Characterisation of the TBEV Spinach2 replicon in mammalian cells. (A) Negative strand RT-PCR analysis in TBEV replicon transfected PS cells. Data are the mean ± SEM of two independent experiments. (B) Culture media collected from TBEV infected PS cells at the indicated times post-infection (pi) were assessed for virus infectivity via plaque assay. Data are the mean ± SEM of three independent experiments. (C-D) Luciferase assays of PS or A549 cells co-transfected with WT TBEV replicon (Hypr or Vs) and pTK-Ren (Renilla) performed at the indicated timepoints post-transfection. Firefly values were normalized to Renilla. Bar heights are the mean ± SEM of two biological replicates. Assays were performed in triplicate. *P=<0.01, **P=<0.001 from WT Hypr determined using a two-tailed Student’s t-test with Welch’s correction.

**Figure 3.**
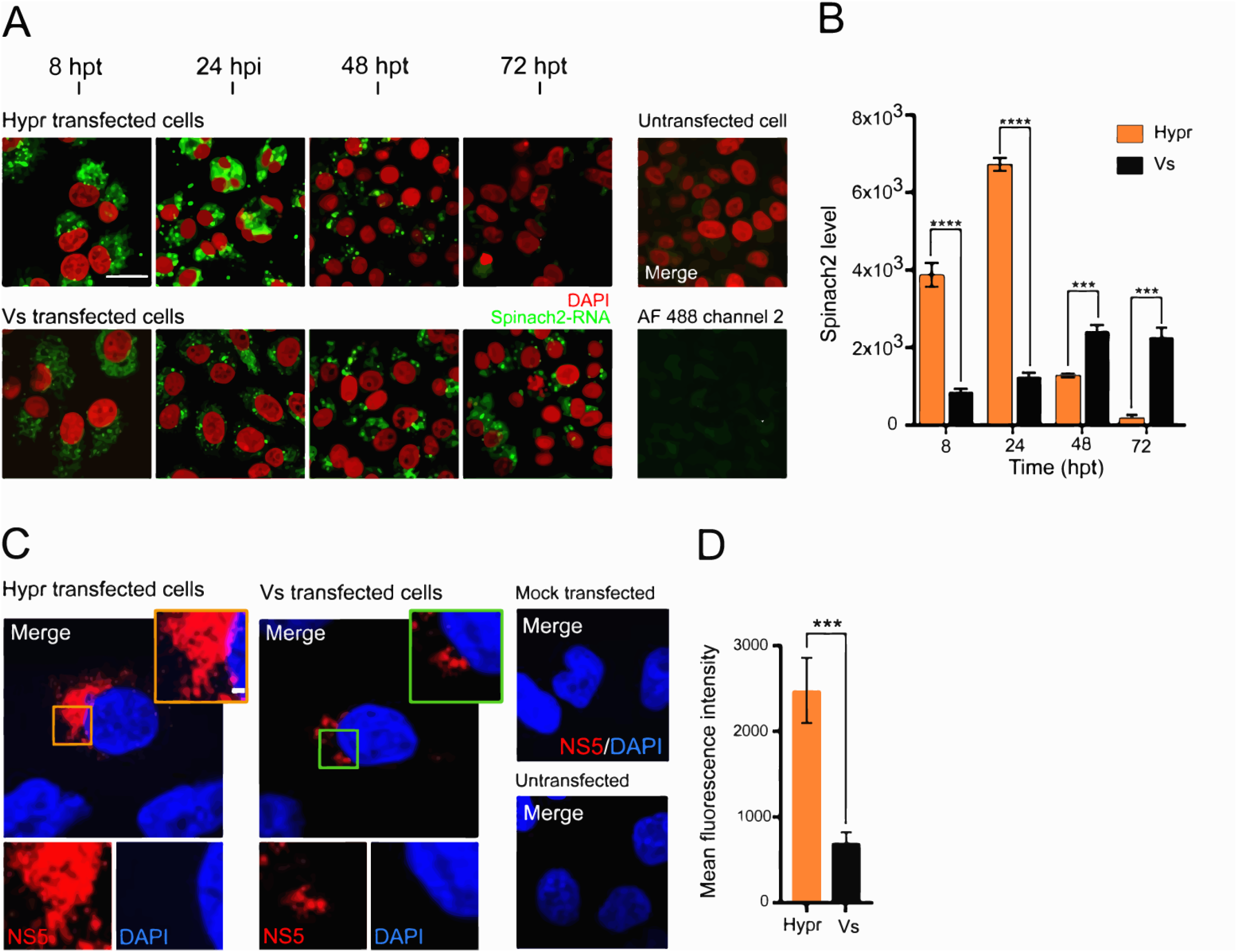
TBEV Spinach2 fluorescence in mammalian cells. (A) Time-course of DFHBI fluorescence (green) in PS cells transfected with Spinach2 tagged Hypr and Vs replicons. Nuclei were stained with Hoechst 33342 (red). Scale bar: 50 μm. (B) Quantification of Spinach2 replicon RNA performed using Fuji (≥ 4 cells). Bar heights are the mean ± SEM of two biological replicates: ***, P=<0.0003, ****, P=<0.0001 from WT Hypr. (C) Subcellular distribution of TBEV NS5 in PS cells transfected with *in vitro* transcribed Hypr and Vs replicons. At 24 hpt, cells were probed using an antiserum specific for TBEV NS5 and stained with goat rabbit IgG (H+L) conjugated to Alexa Fluor^®^ 488 (red). Nuclei were stained with DAPI (blue). Scale bar: 20 μm. (D) Quantification of NS5 expression using Fuji (≥ 4 cells). Bar heights are the mean ± SEM of two biological replicates **, P=<0.001, ***, P=<0.0004 from WT Hypr determined using a two-tailed Student’s *t*-test with Welch’s correction.

**Figure 4.**
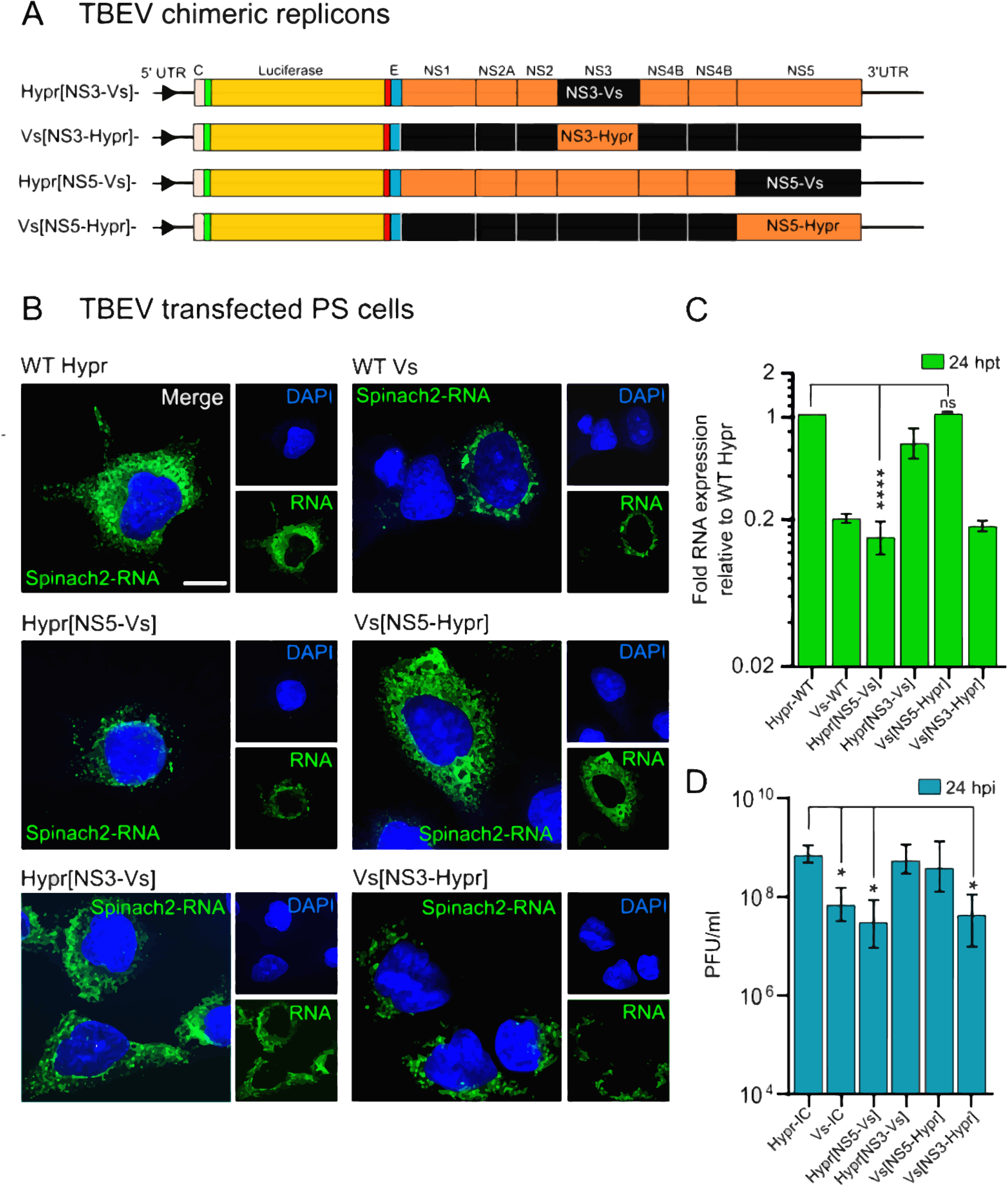
Subcellular localisation of WT and NS5 and NS3 chimeric TBEV replicons in PS cells. (A) Schematic representation of TBEV NS3 (Hypr[NS3-Vs], Vs[NS3-Hypr]), NS5 (Vs[NS5-Hypr] and Vs[NS3-Hypr] chimeras. (B) Replicon transfected cells were treated with DFHBI at 24 hpt and fixed in 4% PFA. Cells were imaged on an Airyscan microscope. Nuclei were stained with DAPI (blue). Scale bar: 10 μm. (C) Quantification of Spinach2-RNA expression of WT *vs* chimeric replicons in PS cells. Bar heights are the mean ± SD of two biological replicates. ****, P=<0.0001 from WT Hypr. ns: no significant difference determined using a one-way ANOVA. (D) Quantification of plaque assays from C2C12 cell supernatants transfected with the indicated TBEV replicons for 24 h. *P<0.01 compared to WT Hypr-IC determined using a one-way Anova. Assays were performed in PC cells.

### Immunofluorescent visualisation of the Spinach TBEV replicon system

To directly visualize replicating TBEV RNA cells were incubated with DFHBI at defined timepoints post-transfection. Confocal analysis showed that the fluorescence of the Spinach labelled RNA was abundant in PS cells transfected with either Hypr or Vs Spinach-replicons, but the cells transfected with the Hypr replicon were characterised by a granular and punctate staining pattern, which peaked at 24 hpt and declined substantially over the next 48 h. In contrast the Vs replicon, showed more diffuse fluorescence, only coalescing into a visible punctate pattern after 48 hpt, which then persisted for the duration of the analysis (Figure 3A). For both Hypr and Vs replicons, the predominant pattern of Spinach fluorescence was perinuclear, most notably at later time points, consistent with the formation of the membranous-web structures frequently observed for positive sense RNA viruses (44). Quantitation of the Spinach fluorescence was consistent with these observations; a spike of Hypr fluorescence was observed followed by a rapid decline, compared to the gradual and more sustained increase detected for Vs replicon (Figure 3B). In addition, replicon expressing cells stained with a rabbit NS5 antisera showed a similar perinuclear aggregation, consistent with this representing the sites of viral replication factories (Figure 3C). The NS5 signal at 24 hpt was significantly greater for Hypr than Vs (Figure 3D). Taken together, these data confirm that Hypr and Vs show variable levels of replication in PS and A549 cells, with a pattern that broadly recapitulates the growth of each virus in mammalian cell culture (31).

### Differential replication of Hypr and Vs strains in mammalian cells is mediated by NS5

The close relatedness of Hypr and Vs offered the opportunity to map the basis of their differential replication pattern through the creation of genetic chimeras between them. In the TBEV replicons described here, the lack of the structural region inferred that the differences between Hypr and Vs observed mapped to the NS coding region, previously shown to dictate virus-mediated CPE in cell culture (31). To further define these differences, hybrid replicons were created through the seamless exchange of Hypr and Vs sequences at the junction of NS3 or NS5, herein termed Hypr[NS3-Vs], Hypr[NS5-Vs], Vs[NS3-Hypr] and Vs[NS5-Hypr] (Figure 4A). When PS cells were transfected with each chimeric replicon and assayed for Spinach fluorescence, the prevalent early fluorescence signal associated with the Hypr replicon was lost upon exchange of the Hypr NS5 region with that of Vs. Conversely, the weaker fluorescence associated with the parental Vs replicon increased substantially when the Vs-NS5 was replaced with that of Hypr. In contrast, exchange of the NS3 coding region had little effect (Figure 4B), with measurements at 24 hpt showing that the fluorescence associated with the Vs background carrying only the Hypr-NS5 sequence was as high as the parental Hypr strain, whilst that of Vs carrying the Hypr-NS3 sequence showed no significant difference (Figure 4C-D). Thus, the replication phenotypes observed for Hypr and Vs, which match the differences seen in CPE detected for the parental viruses, lie within the NS5 region of the TBEV genome.

TBEV is an arbovirus that replicates naturally in both tick and mammalian cells. To address if the differences observed in the replication kinetics of Hypr and Vs were conserved in invertebrate cells, *Spodoptera frugiperda* (Lepidoptera) cells were transfected with the parental and chimeric replicons and the Spinach associated fluorescence signal assessed. In contrast to that observed in mammalian (PS and A549) cells, the Spinach related fluorescence intensities of the parental Hypr and Vs replicons showed no obvious differences when imaged using standard confocal microscopy (Figure 5A, upper panel) or gated stimulated emission depletion (gSTED) microscopy, with neither the number nor the size of the fluorescent foci varying (Figure 5A, lower panels). Similarly the NS3-NS5 chimeras showed no differential signal (Figure 5B). In all cases, the fluorescent signals were diffuse throughout the cell, which contrasted with the perinuclear distribution observed in PS cells. Quantification of the Spinach fluorescent signal at 24 hpt in four representative cells from each transfection showed no significant differences amongst the Hypr, Vs or NS3-NS5 hybrid replicons, indicating comparable levels of replication (Figure 5C). These data suggest that the differences between Hypr and Vs in mammalian cells, which maps to the NS5 region of the TBEV genome, is a product of the mammalian cell environment and not a product of variable RdRp activity associated with each strain.

**Figure 5.**
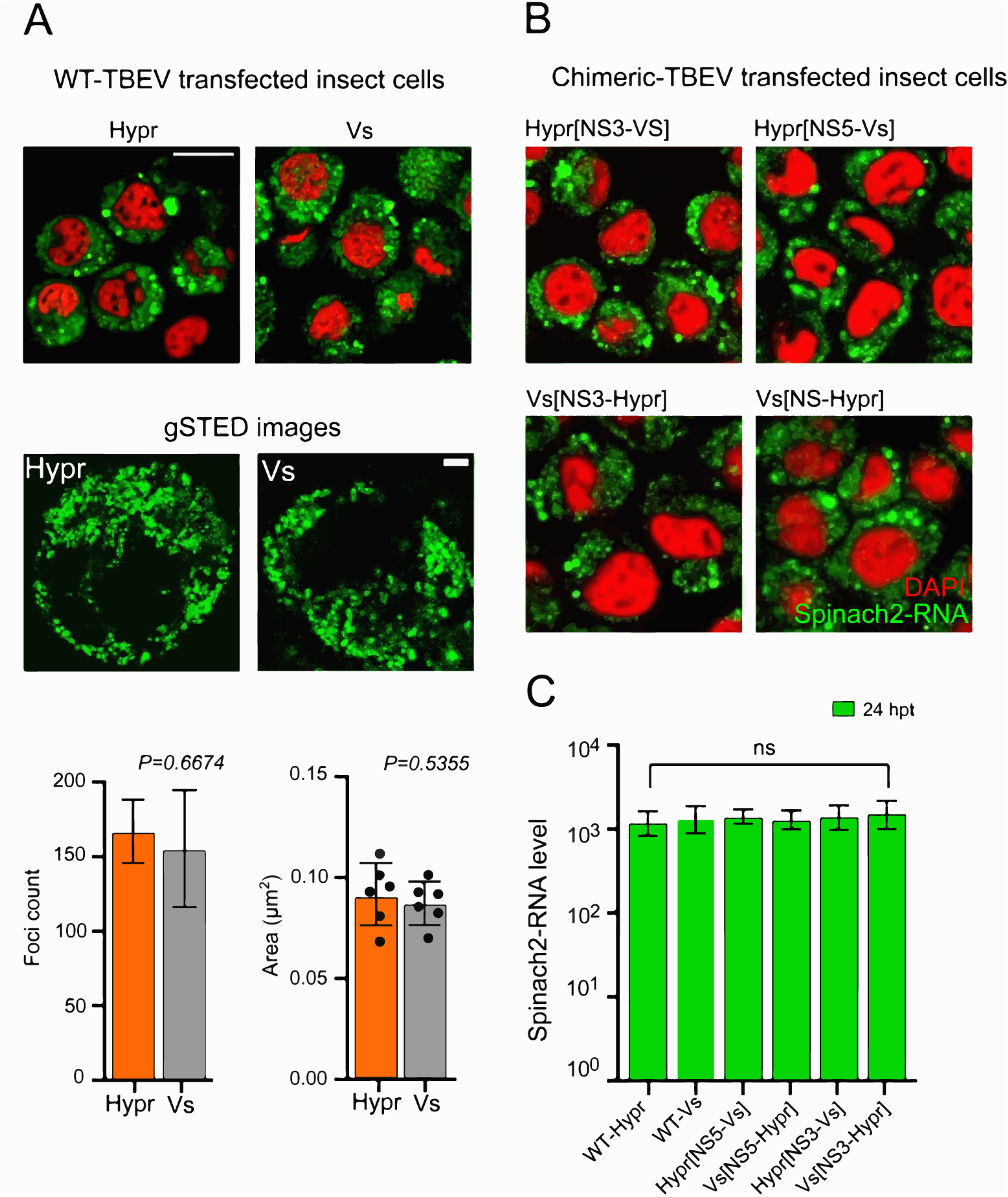
Subcellular localisation of TBEV WT and NS5 and NS3 chimeric replicons in insect cells. (A) Sf9 cells were transfected with SP6 *in vitro* transcribed Spinach2 tagged (A) WT Hypr, Vs or (B) chimeric replicons. (A) At 24 hpt, cells were incubated with DFHBI and live cell imaged on a Zeiss LSM780 confocal or Leica TCS SP8 gSTED microscope. Scale bars: 50 μm and 5 μm, respectively. Spatial data for gSTED imaged Spinach2-RNAs determined using Fuji (≥ 6 cells). Data were used to determine the Spinach2-RNA foci count per cell and average size per focus for Hypr and Vs (lower graphs). (B) Spinach fluorescence from cells transfected with the indicated hybrid replicons (green). Cells were co-stained with DAPI (blue). Scale bar: 20 and 5 μm, respectively. (C) Quantification of WT and hybrid Spinach2 RNA expression using Fuji (≥ 4 cells for each hybrid). Bar heights are the mean ± SEM of three biological replicates. ns: no significant difference from WT determined using a one-way ANOVA.

### Hypr and Vs differentially induce the innate immune response

In insect cells, the interaction of TBEV with the innate immune system plays a role in the outcome of infection. The major mechanism of antiviral defence includes the siRNA systems, which responds to viral RNAs to supress viral replication through the Dicer and Argonaut pathways (45). In mammalian cells, the detection of viral RNAs by sensors of the innate immunity system triggers a signalling cascade that results in the induction of interferon response and in some cases, cell death (41). To investigate whether immune sensing mediates the differences observed between Hypr and Vs strains in mammalian cells, the level of key host defence markers were compared between replicons. These included: TIAR and G3BP1, markers for stress granule formation that have previously been shown to bind to the TBEV genome (46, 47); phosphorylated AKT, a pro-survival kinase activated in TBEV infected cells (48); IRF3, the canonical interferon response factor previously shown to be induced by TBEV (17), and caspases 8 and 3, inducer and executioner caspases respectively of the apoptosis cascade (49). Upon analysis, stress granule formation was more extensive in Vs compared to Hypr replicon cells (Figure 6A) and, when co-stained for G3BP1 and dsRNA as a marker for the TBEV replicative intermediate, significant colocalization was observed in Vs but not Hypr transfected cells (Figure 6B, left panel). Quantitative analysis showed that the percentage of dsRNA fluorescence that co-localised with G3BP1 in Vs transfected cells was significantly elevated compared to Hypr where the majority of dsRNA did not associate with G3BP1 (Figure 6B, right panel). Similarly, the level of phosphorylated AKT (Serine 437) in Vs replicon transfected cells was 5-fold higher than was observed in Hypr transfected cells (Figure 7A). Induction of the interferon response measured through the levels of phosphorylated (Serine 396) and non-phosphorylated IRF-3 showed a reciprocal pattern, with only low levels of pIRF-3 observed in Vs replicon cells compared to Hypr (Figure 7B). Apoptosis was also differentially induced between the two TBEV strains; Hypr replicons showed high levels of activated caspase-3 and caspase-8 compared to the low levels observed for Vs (Figure 8). Taken together, these data suggest that differential host cell defence responses to Hypr and Vs strains contribute to the replication characteristics observed for these viruses in mammalian culture systems.

**Figure 6.**
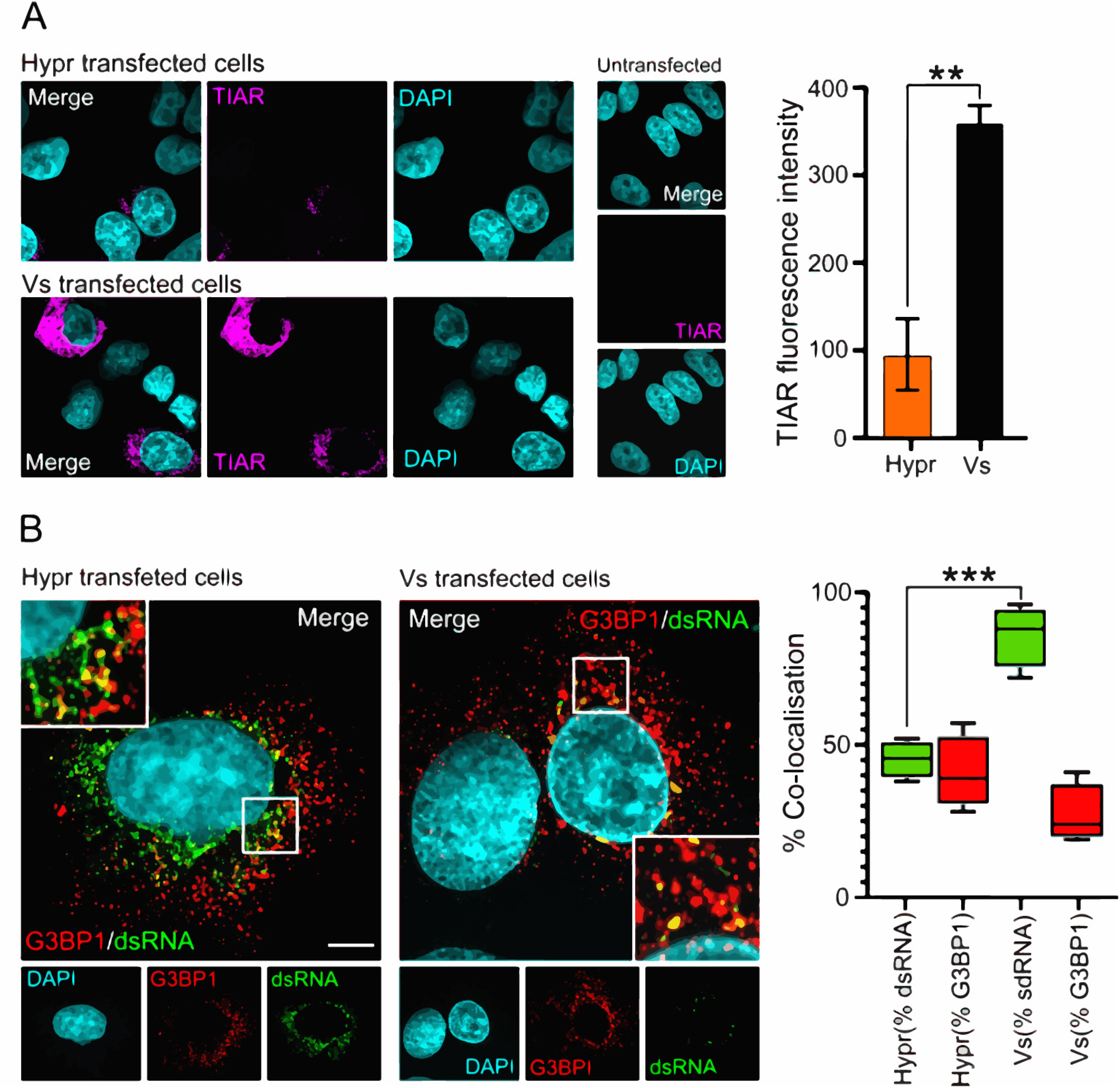
Formation of stress granules in TBEV replicon cells. (A) Replicon and mock transfected cells were fixed with 4% PFA at 24 hpt and stained with anti-TIAR (magenta) and DAPI (cyan). Cells were imaged on a Zeiss LSM780 confocal microscope. Scale bar: 20 μm (left panel). Quantification of TIAR levels in Hypr and Vs transfected cells (right panel). Bar heights are the mean ± SEM of two biological replicates. (B) Transfected cells were stained with anti-G3BP1 (red), J2 anti-dsRNA (green) and DAPI (cyan) and imaged on an Airyscan microscope. Scale bar: 10 μm or 1 μm, respectively (left panel). Right panel of B indicates quantification of the colocalization of dsRNA with G3BP1 (green) or G3BPI colocalised with dsRNA (red). Manders’ overlap colocalization coefficients were calculated from ≥10 cells using Fuji, from at least two independent experiments. **, P=<0.001, ***, P=<0.0001 from WT Hypr determined using a two-tailed Student’s *t*-test with Welch’s correction.

**Figure 7.**
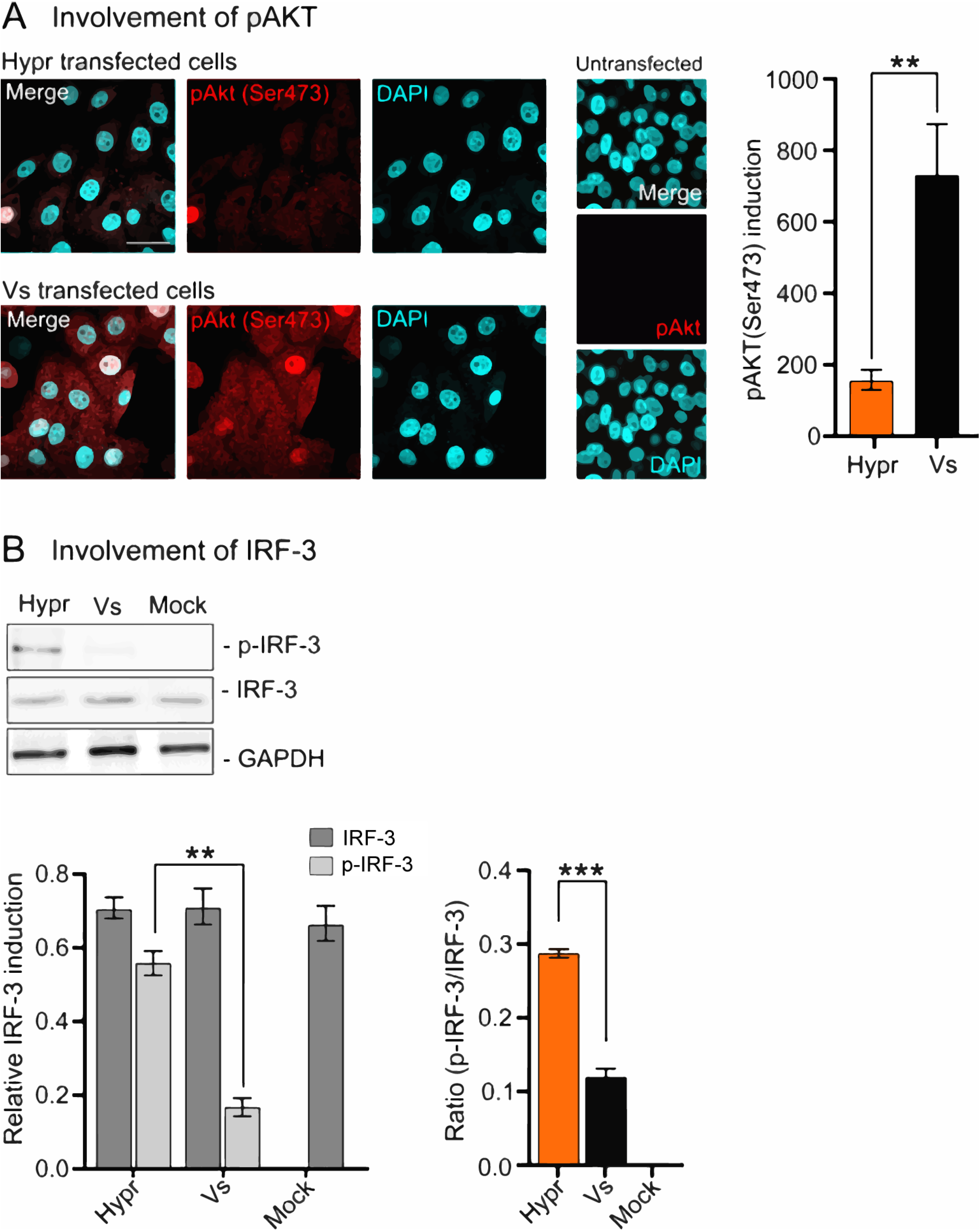
Phospho-AKT (Ser473) and IRF-3 levels in TBEV replicon cells. (A) Immunofluorescence analysis of pAKT (Ser473) in Hypr and Vs transfected PS cells at 24 hpt. Cells were fixed with 4% PFA and stained with anti-pAKT (Ser473) (red) and DAPI (cyan), and imaged on a Zeiss LSM780 confocal microscope. Scale bar: 50 μm (left panel). Quantification of pAKT (Ser473) in Hypr and Vs replicon transfected cells (right panel). Bar heights are the mean ± SEM of two biological repeats. (B) IRF-3 status in replicon transfected cells at 20 hpt analysed by western blot for anti-pIRF-3(Ser396), and anti-IRF-3. Lower panels represent the quantification of p-IRF-3(Ser396), total IRF-3 and the ratio of p-IRF-3 to total IRF-3. Relative expression was normalised to GAPDH. Bar heights are the mean ± SEM of three biological replicates. **, P=<0. 0.0095, ***, P=<0.0001 from WT Hypr determined using a two-tailed Student’s *t*-test with Welch’s correction.

**Figure 8.**
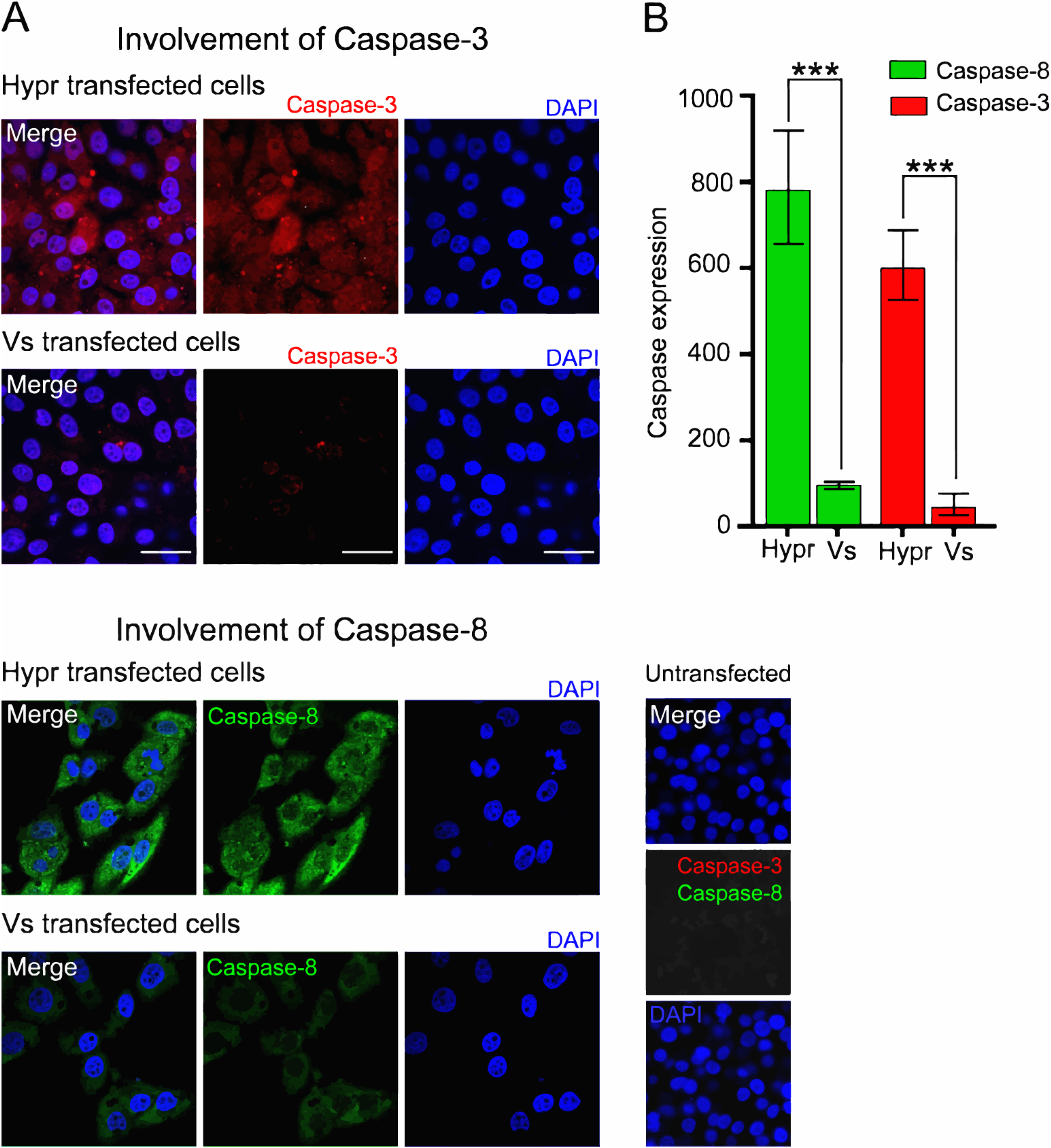
Hypr and Vs TBEV strains differentially induce caspase-3 and −8. (A) Immunofluorescence analysis of caspase −3 and −8 in Hypr and Vs transfected PS cells at 16 hpt. Cells were fixed with 4% PFA, stained with anti-Caspase-3 (red), anti-Caspase-8 (green) and DAPI (blue) and imaged on a Zeiss LSM780 confocal microscope. Scale bar: 50 μm. (B) Quantification of Caspase-3 and −8 staining determined using Fiji. Bar heights are the mean ± SEM of three biological replicates. ***, P=<0.0001 from WT Hypr determined using a two-tailed Student’s *t*-test with Welch’s correction.

## Discussion

TBEV infection continues to spread throughout Europe with emergence now reported in previously unaffected areas. In infected patients, specific TBEV strains lead to diverse clinical symptoms, ranging from mild to severe, also mirrored by their cytopathic effects (CPE) in culture (19, 20, 32). The mechanisms underlying the variability of these clinical symptoms remain unknown but are likely mediated by the characteristics of the host and specific features of the TBEV genome (50). In nature, TBEV strains with both higher (strain Hypr) and lower (strain Vs) pathogenicity have been described. We therefore constructed Spinach2-tagged replicons and chimeras of Hypr and Vs to investigate the basis of their differential pathogenicity. Previous TBEV replicons have expressed eGFP or luciferase, which reports only the final steady state level of viral protein expression, as opposed to the direct measurement of viral RNA accumulation and the localisation of viral RNA which the Spinach-tagged system permits (32).

Using these replicons, we show that the Hypr strain replicates to high levels at early time points peaking at 24 hpt and declining thereafter (Figure 1–4). By contrast, the Vs replicon produces lower and more sustained levels of viral RNA that persist for longer (Figure 1–4). These phenotypes were confirmed by assessment of Hypr and Vs viruses in two independent mammalian cell lines, validating the replicons for further chimeric assessments of Hypr and Vs pathogenicity. Through construction of the Hypr/Vs hybrids (Figure 4), we show that the key determinant of differential virus replication kinetics maps to the NS5 region of the genome, with exchange of the NS3 region leading to no discernible changes in Hypr or Vs replication. Of note, the differences in Hypr and Vs systems were restricted to mammalian cell culture systems, with comparable replication kinetics observed among all Hypr and Vs replicons in invertebrate cells (Figure 5). This strongly implicates the host cell response induced by NS5 as opposed to its RNA-dependent RNA polymerase (RdRp) activity, as the key mediator of the differences between Hypr and Vs strains. NS5 possesses RNA cap methyltransferase (MTase) and RdRP activity which mediates TBEV replication. Alignment of the ~911 amino acid sequences of Hypr and Vs NS5 shows 50 changes, 16 of which lie within the N-terminal O-methyltransferase (O-MT) domain, whilst 34 map to the larger RdRp, a small mutational bias towards O-MT. NS5 has been ascribed a range of functions in TBEV infected cells in addition to its direct role in virus replication, most notably its influence on Jak-STAT signalling (22, 24) and RANTES induction through the activation of IRF-3 signalling that is dependent on RIG-I/MDA5 (17). Of note, we observed several phenotypic differences in cells expressing Hypr and Vs Spinach-replicons, including stress granule formation (Figure 6), AKT and IRF3 activation (Figure 7), and caspase 8- and 3- induction (Figure 8). Antiviral innate immune responses such as stress granule formation, observed for the Vs replicon, block translation and arrest viral gene expression (46, 49), which may explain its reduced level of replication when compared to Hypr. In addition, Hypr-NS5 induced IRF3 expression would be predicted to drive innate immunity and subsequent neuroinflammation, a common feature of pathogenic TBEV viruses. Recent studies also highlight how cell type-specific innate immunity contributes to shaping TBEV tropism and ultimately disease outcome in human brain cells (7).

By extension, differential innate responses might be critical to the clinical outcome of flaviviruses associated with a broad disease spectrum (51), as is the case for Vesicular Stomatitis Virus, which can switch tropism from the peripheral to the central nervous system based on the strength of the IFN response in the lymph node subcapsular region, which acts as a gateway between the vasculature and the CNS (52, 53). Minor changes in innate immune antagonism due to virus emergence from the quasispecies may thus translate to apparent gross clinical differences observed across the TBEV strains. It is notable that differential triggering of innate sensors such as RIG-I also underlie the differential cytokine induction observed during Influenza virus infection, also typified by major differences in clinical outcome depending on the virus isolate (54).

No specific treatments for TBEV infection are currently available and control is achieved by a prophylactic vaccine offered in endemic areas and for those who travel to them (7). Our data suggest that interventions to moderate the activities of NS5 may reduce the virulence of TBEV through the suppression of its ability to modulate host defences. The finding that the replication kinetics of specific TBEV strains can be easily monitored by the Spinach-replicon system could allow screening of a range of current and emerging TBEV strains as part of preparedness measures. The importance of key amino acid substitutions in pathogenic vs non-pathogenic TBEV with such replicons can also bypass studies with full-length infectious cDNA clones that require biosafety level 3 containment, advancing our understanding of TBEV pathogenesis. A comprehensive assessment of these replicons in a range of neuronal and tick cell systems will form the basis of future studies that aim to fully define how NS5 and innate immune defences control TBEV pathogenicity and the subsequent variability in disease presentation.

## Acknowledgments

We respectfully acknowledge the late Tamara Gritsun for her dedication to studies of TBEV, careful supervision and for the original clones that permitted this study. For part of the work NG was the recipient of a Dean’s Research Award from the University of Leeds. We are grateful to Michelle Peckham (University of Leeds) for access to the Zeiss LSM880 Airyscan confocal microscope (funded by Wellcome Trust grant 104918/Z/14/Z), the staff of the Micron Facility, (Biochemistry Department, Oxford University), and the STFC Central Laser Facility, Culham for access to the gSTED microscopy suite. We also thank Carsten Zothner for technical assistance, Holly Bratcher (University of Oxford) and Benjamin Neuman (Texas A&M University-Texarkana) for advice.

